# Dissection of Catalytic Site in Crucial Gut Microbiome Enzyme: Bile Salt Hydrolase

**DOI:** 10.1101/714493

**Authors:** Yashpal Yadav, Mrityunjay K. Tiwari, Deepak Chand, Debjoyati Boral, Archana Pundle, Sureshkumar Ramasamy

## Abstract

Bile Salt Hydrolases (BSHs) are enzymes from enteric bacteria that catalyze the hydrolysis of Bile Acids and consequently promote the reduction of cholesterol level in the mammalian body. Out of several reported BSHs, the *Enterococcus faecalis* BSH (*Ef*BSH) has been reported to have the highest enzymatic activity. Herein, we have investigated the mechanistic details of the *Ef*BSH activity. The study was carried out employing two mutants of *Ef*BSH: E269A and R207A, which shows differential catalytic activity. The mutant E269A exhibits significant loss in the BSH activity with an increased affinity towards the substrate as compared to R207A mutant. Further, R207A was found to be involved in allostery with an increased *Ef*BSH activity towards tauro-conjugated bile acids. The structural and electrostatic force analyses of the active sites of the E269A mutant and the wild type *Ef*BSH (wt *Ef*BSH) revealed that the interaction between Glu21 and Arg207 is the determining factor in maintaining the dynamic allostery and high activity of *Ef*BSH.

## Introduction

Bile Salt Hydrolase (BSH) is a class of enzymes that catalyze the hydrolysis of the amide bonds of Bile Acids (BA’s) (Duggleby et al., 1995). The deconjugated bile acids, the bile acids obtained after hydrolysis, have lower affinity towards the bile acid transporters such as ASBT and NTCP, which lead to reduction in the rate of enterohepatic recirculation (Hu, Iwata, Cameron, & Drew, 2011; Zhou et al., 2014), and hence results into excretion of bile acids through feces (Chand et al., 2017). Begley in 2006 reported the bile salt hydrolase activity as one of the probiotic selection criteria and a desirable trait for the survival of microflora in the human gut (Begley, Hill, & Gahan, 2006). Recent studies also suggest that the growth of BSH positive organisms is better in the presence of bile acids (Jones, Begley, Hill, Gahan, & Marchesi, 2008). The microbial BSH activity plays a crucial role in bile acid homeostasis and maintenance of bile acid pool. Hence deconjugation of bile acid in the gut is of paramount importance in context to cholesterol reduction.

Structural analysis of bile salt hydrolases revealed the active site pocket, which is constituted of four distinct loops (denoted as loop1-loop4) and an assembly loop or tetramer loop (Fig. 1a & b) that holds the “dimer of dimer” to form a tetramer (R. S. Kumar et al., 2006; Rossocha, Schultz-Heienbrok, von Moeller, Coleman, & Saenger, 2005). Loop1, loop2 and loop3 allowed the entry of substrates into the active site of the enzyme and referred as ‘Site A’, whereas the site of glycine release is referred to as Site L as shown below (Fig. 1b) (Panigrahi, Sule, Sharma, Ramasamy, & Suresh, 2014; Rossocha et al., 2005). With few exceptions, most of the gram positive and gram negative organisms possess the tetrameric form of BSH. Distinct clustering of gram positive and gram negative bacterial BSH were observed owing to indel mutation (20-22 amino acids) in the tetramer loop region (Panigrahi et al., 2014). Theoretical prediction states that the presence of tetramer loop provides thermostability to gram positive BSHs as compared to gram-negative BSHs (Panigrahi et al., 2014). Recently, Bile Salt Hydrolase from *Enterococcus faecalis* (*Ef*BSH) showed several folds higher specific activity than that for the previously reported BSHs (Chand, Ramasamy, & Suresh, 2016). This enzyme acts on both the *tauro-* and *glyco-*conjugated bile acids. Further, the structural studies of *Ef*BSH report that the binding pocket of this enzyme has more hydrophobic residues, particularly aromatic residues, near site A (Fig. 1b) that consists of loop1 and loop2 and where the cholyl moiety binds (Chand, Panigrahi, Varshney, Ramasamy, & Suresh, 2018; Panigrahi et al., 2014). Also, RMSD calculation of individual monomer unit of tetrameric *Ef*BSH exhibited high RMSD, indicating the allosteric nature of this enzyme (Chand et al., 2018). Moreover, disruption of loop2 is believed to transmit allosteric effects to remaining chains *via* tetramer loop (assembly loop) and the residues of loop 1 and loop2 *viz*. F18, Y20, and Y65 align the substrate in an orientation that has been proposed to account for the higher specific activity by forming the better catalytic framework in the *Ef*BSH (Chand et al., 2018). However, the detailed basis for the higher activity of the enzyme and its allosteric nature is still obscure. Owing to the pharmacological importance of BSH enzymes, it is therefore necessary to understand the structure and activity correlation of these enzymes. Furthermore, pinpointing the residues responsible for the increased activity may help to tailor the molecule to obtain enhanced activity.

**Fig. 1:**
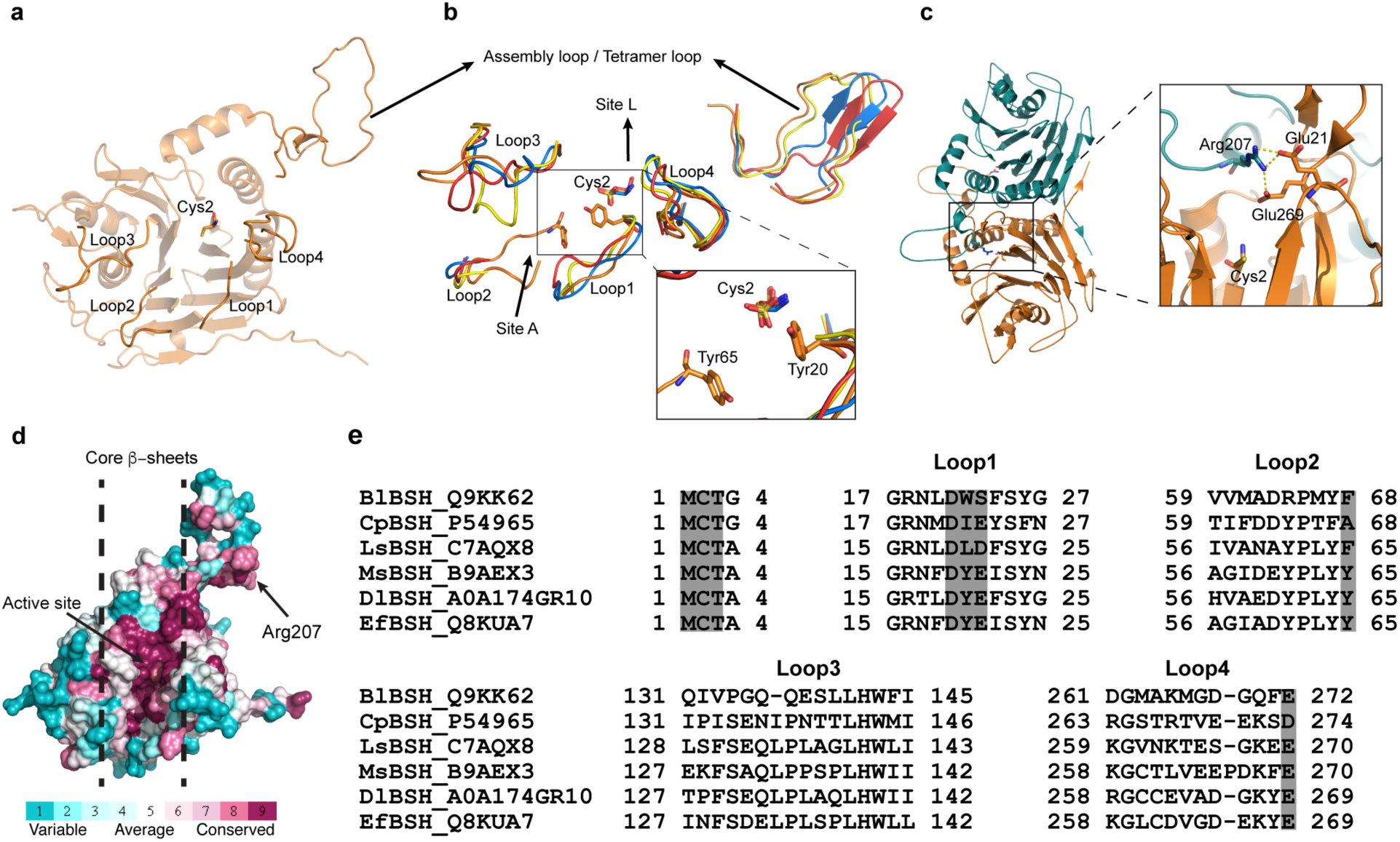
Overall structure of Bile Salt Hydrolase (BSH): **(a)** shows the arrangement of loop1 to loop 4 near active site and tetramer loop or assembly loop with reference to whole molecule of wild type *Ef*BSH. **(b)** Superposed image of reported BSH structure i.e. *Ef*BSH (orange), 4wl3; *Ls*BSH (yellow), 5hke; *Bl*BSH (Red), 2hez; and *Cp*BSH (Blue), 2rlc; showing the entry and exit site of substrates. Inset image shows two tyrosine residues reported to align substrates. **(c)** Dimeric form of wt*Ef*BSH revealing the Arg207 of chainB protrudes above the active site of chainA. Inset image displays Arg207 from chain B protruding inside the active site pocket and hydrogen bonded to residues of chain A. **(d)** Consurf image showing conservation of residues in core β-sheets and assembly loop. The conservation score is color ramped from cyan (variable) to white (average) to red (conserved). **(e) A**lignment of bile salt hydrolases from different species and comparision of loop1 to loop4. Grey highlighted region showing the differences in the residues in loop region.

Herein, we report the structure of two mutants, *Ef*BSH E269A (E269A) and *Ef*BSH R207A (R207A) and their biochemical characterization, which has provided a detailed understanding of the higher catalytic efficiency and allostery. Apart from loop1 and loop2, loop4 also plays crucial role in the activity of *Ef*BSH possessing the E269 residue which forms H-bond with core substrate binding residue Glu21 and maintains the active site geometry for the better catalytic framework. Consurf analysis (Ashkenazy et al., 2016) showed the tetramer loop and α-helices to be highly variable, whereas the core β-sheets are highly conserved (Fig. 1d). Structural analysis showed that Arg207 in tetramer loop protrudes into the active site of an adjacent subunit (Fig. 1c), which influences the active site geometry. Hydrogen bonding network showed that Arg207 interacts with two residues *viz*. Glu269 and Glu21 of adjacent subunit (Fig. 1c inset). However, levels of this interaction for each monomeric chain are found to be different. In perspective, we propose here, the open and closed conformation of BSH active site owing to the interaction of Arg207 with Asp21 that controls the movement of Tyr20, which acts as a gatekeeper to Cys2 catalytic residue near the active site.

## Materials and Methods

All the chemicals are procured from Sigma (Sigma, USA) and Merck (Merck, USA). Media components are obtained from Hi-Media, plasmid isolation kits from Qiagen (Germany), Quick Change site-directed mutagenesis kit from Stratagene (USA), restriction enzymes and ligases from New England Biolabs (NEB, UK), Oligonucleotide DNA from Eurofins (India) and pET 22b(+) vector and Nova Blue competent cells from Novagen (Novagen, USA). The *Ef*BSH wild type clone was obtained as a gift from Dr. Deepak Chand (Chand et al., 2016).

### Site directed mutagenesis

The mutants of *Ef*BSH *viz*. R207A, E269A, E269Q and E269D were prepared using the wild-type *Ef*BSH plasmid. The primer sequences used for creating mutations are listed in supplementary information Table 2. The PCR was performed using Thermal cycler from AB sciences (USA), at 95 °C for 3 min and 18 cycles (94/ 30 sec, 55 °C for 30 sec, 72 °C for 6 min 30 sec.) and final ramping of 72 °C for 10 min. The PCR mixture was then treated with *Dpn*I enzyme to degrade the methylated parental DNA and then incubated at 37 °C for 3 hrs. *E*.*coli* DH5α cells were transformed with 5µl of PCR mixture. The plasmids were purified and sequenced to confirm the desired mutation.

### Heterologous expression and purification of *Ef*BSH and the mutant proteins

Recombinant plasmid containing the wild-type *Ef*BSH and R207A, E269A, E269Q and E269D mutants were transformed into the expression host *E. coli* BL21 (DE3) cells. The wild-type *Ef*BSH and mutants were purified using the protocol reported in our previous studies (Chand et al., 2016). The cell pellet was resuspended in the lysis buffer composed of 25mM Tris-Cl pH 8.0, 500mM NaCl, 0.1% (v/v) Triton X-100 and 1 mM DTT. The cell pellets were then sonicated (E-Squire Biotech, India) and purified in two steps: first by employing Ni-NTA affinity chromatography followed by Size Exclusion Chromatography (SEC) using Superdex 200 (30/300 GL). The SEC was performed with 20 Mm Tris-Cl pH 7.5 buffer containing 150 mMNaCl to purify the protein to homogeneity. The purity of protein was assessed by SDS-PAGE. The fractions from gel filtration were pooled and concentrated to 10 mg/ml for crystal set-up and for biochemical as well as kinetic studies.

### Measurement of Enzyme Activity and Kinetic parameter of *Ef*BSH mutants

Bile Salt Hydrolase activity was determined by Ninhydrin assay (Chand et al., 2016; R. S. Kumar et al., 2006). 50 µl mixture of enzyme and GCA incubated at 37 °C for 10 min and then the reaction was quenched using an equal volume of 15% v/v TCA (Trichloro Acetic Acid). The enzyme units were determined in terms of glycine release. One unit is defined as the amount of enzyme that liberates 1µmol of glycine per minute. The kinetic constants, K_m_ and V_max_, of *Ef*BSH mutants were calculated using varying substrate concentration ranging from 0.5 mM to 60 mM in triplicates using GraphPad Prism version 5.0 (Graphpad Inc., USA). GCA was used as a substrate (Sigma-Aldrich, St. Louis, MO, USA) for calculating the kinetic parameters. Kinetic parameters were calculated using nonlinear regression by fitting the kinetic data.

#### (iv) Crystallization, diffraction and data collection

The purified homogenous protein was concentrated to 10 mg/ml using Amicon ultra-15 ml 30 kDa cut-off centrifugal filters. Preliminary crystal hits were obtained from commercial screens such as PEG Ion, Index, Nextal Pact, JCSG plus. Screening was performed using vapor diffusion method in 96 well MRC2 sitting drop plates (Hampton) having 200 nl of both protein and screening solution. Crystals of E269A mutant was obtained in JCSG plus screen comprising 0.14 M CaCl2, 0.07M CH3COONa, pH 4.6 and isopropanol 14% v/v. R207A mutant crystallized in 0.1M Sodium Citrate Tribasic Dihydrate, pH 5.5 and 22% PEG1000. The crystals were flash frozen in liquid nitrogen (100 K) without cryoprotectant and data were collected at PXBL-21 (Indus-2), RRCAT, Indore, India (A. Kumar et al., 2016).

### Structure determination and refinement

The data sets were recorded with a Rayonix MX225 CCD imaging plate detector system. The data were processed in iMosflm (Battye et al., 2011) and scaled in the AIMLESS programs. 5% of the obtained data was reserved for cross-validation. Details of data collection statistics is provided in Table 1. The structure was determined using the molecular replacement method using data between 50 and 2.0 Å in the Phaser-MR (McCoy et al., 2007). The crystal structure of BSH from *Enterococcus faecalis* (PDB ID: 4wl3, chain A) was used as a template for the search model. Model building and structure refinement were done iteratively using Coot (Emsley & Cowtan, 2004) and Refmac5 (Murshudov et al., 2011) (CCP4 suite) till R-free decreases to an allowed range. Water molecules were manually added in coot at peaks of density above 4σ in the asymmetric unit. Finally, the stereochemistry of the structure was validated by the PROCHECK program. All the structures were made using Pymol.

**Table 1:** X-ray diffraction and Data collection statistics for *Ef*BSH mutant structures viz. R207A and E269A. Numbers in parentheses correspond to highest resolution shell.

|  | **R207A_mutant** | **E269A_mutant** |
| --- | --- | --- |
| **Data Statistics** |  |  |
| Space group | P2<sub>1</sub> | P2<sub>1</sub> |
| X-ray source | INDUS-II, RRCAT | INDUS-II, RRCAT |
| Wavelength | 0.97 | 0.97 |
| Resolution (Å) | 38.97—2.0 | 41.14-2.10 |
| Unit cell parameter | a=59.33, b=156.45, c=72.91;<br>$\alpha=\gamma=90$,<br>$\beta=98.62$ | a=59.84, b=156.32, c=73.15;<br>$\alpha=\gamma=90$,<br>$\beta=98.8$ |
| Molecule per asymmetric unit | 4 | 4 |
| Matthews Coefficient (Å<sup>3</sup> Da<sup>-1</sup>) | 2.21 | 2.20 |
| Solvent content (%) | 44.38 | 44.0 |
| Total No. of reflection | 335415(17397) | 290807(17176) |
| No. of unique reflection | 83294(4365) | 77248(4592) |
| Multiplicity | 4.0 (4.0) | 12.0 (7.4) |
| Completeness (%) | 98.7 (96.5) | 100%(100%) |
| Average I/$\sigma$(I) | 5.0 (2.0) | 3.8(3.7) |
| Rsym or Rmerge (%) | 5.6 (0.41) | 6.1 (0.74) |
| **Refinement Statistics** |  |  |
| Rfactor (%) | 20.73 | 20.35 |
| Rfree (%) | 23.62 | 24.8 |
| RMS Bond Length (Å) | 0.0133 | 0.0152 |
| RMS Bond Angle (°) | 1.6380 | 1.7650 |
| Overall B (Isotropic) from Wilson Plot | 13.4 | 13.3 |
| Most favourable region (%) | 96.6 | 96.2 |
| Additional allowed Region (%) | 3.0 | 3.2 |
| Outlier region (%) | 0.5 | 0.6 |
| Disallowed region (%) |  | 0 |

**Table 2:** Electrostatic Force of binding in pN for Arg-Glu zwitterion pair for the constrained optimized geometries of the active site of *Ef*BSH at the M06-2X(water)/6-31G** level of theory.

| Electrostatic Force of binding for Arg-Glu zwitterion pair |  |  |  |  |  |  |  |  |
| --- | --- | --- | --- | --- | --- | --- | --- | --- |
|  | Chain A |  | Chain B |  | Chain C |  | Chain D |  |
|  | WT | E269A | WT | E269A | WT | E269A | WT | E269A |
| Mulliken | -31.8 | -43.8 | -29.9 | -41.3 | 1.0 | -47.9 | -35.4 | -39.6 |
| NBO | -44.0 | -60.9 | -32.6 | -57.9 | 13.9 | -66.0 | -51.2 | -54.4 |

WT= wild type
E269A = E269A mutant type
Mulliken = Forces involving Mulliken Charges
NBO = Forces involving NPA charges

### Computational Details

All the DFT calculations until unless mentioned specifically were carried out at the M06-2X/6-31G** level of theory (Zhao & Truhlar, 2006, 2008) using the Gaussian 09 suite of quantum-chemical programs (Gaussian09, 2009). The solvent effect is introduced into the calculations employing Conductor like Polarization Continuum Model (CPCM) considering water as the solvent (ε = 78.3553) (Barone & Cossi, 1998; Cossi, Rega, Scalmani, & Barone, 2003).

The amino acids constituting the active sites in all the four chains (A-D) of the wild-type and the E269A mutant are chopped out from the crystal structure geometries of the two proteins. The N-terminals of all the amino acids (other than cysteine, which is the N-terminal amino acid in all chains) were capped with the acyl group and the C-terminals were capped with N-methyl group preserving the coordinates of all the heavy atoms (atoms other than hydrogen) to their crystallographic positions. Hydrogen atoms were added to the resultant structures in order to complete the valency of all the atoms present in these structures. The geometries were optimized under the constrained condition where coordinates of all the heavy atoms were fixed at their crystallographic positions. Thus, only externally added hydrogen atoms were allowed to reorient its positions during the optimization in order to keep the active site structures preserved to their crystallographic form. Such strategy is a common practice in modeling the enzymatic reactions employing quantum cluster approach where coordinates of some atoms are locked to their crystallographic positions in order to preserve the basic architecture of the active site pocket (Himo, 2017). The coordinates of all the geometries obtained after the constrained optimization are provided in the supporting information.

S-H bond cleavage of the N-terminal cysteine residue is reported to be one of the key steps in the catalytic activity of the BSHs (Lodola et al., 2012; Oinonen & Rouvinen, 2000). The electrostatic influence of all the atoms (present in the active site only) on the S-H bond cleavage is evaluated in order to estimate the influence of the E269A mutation on the activity of the *Ef*BSH, as a representative case for all the steps involved in the bile acid hydrolysis. To evaluate this effect, the electrostatic force analysis approach is employed. The electrostatic forces are calculated employing Coulomb’s law considering long range electrostatic influence of every atom present in the active site region are significant. Atoms in the molecules are considered as point charges. The charges are obtained from the quantum chemical calculations. The NBO (Reed, Weinstock, & Weinhold, 1985) and Mulliken (Mulliken, 1955) charges are used to calculate the electrostatic forces. The employed electrostatic force approach has shown to give a reliable quantitative trend in binding among atoms as indicated in our previous studies (Tiwari, science, & 2017, n.d.).

To calculate the electrostatic influence of the non-cysteine atoms of the active site at the S-H bond cleavage, the cysteine molecule (in dipeptide form here) is considered to be made up of two fragments: (i) the hydrogen atom (fragment 1) and (ii) rest of the part (fragment 2). The contribution of non-cysteine atoms on the S-H bond cleavage is calculated employing the following procedure. Every non-cysteine atom is considered to be interacting with each atom of the cysteine from both the fragments. A component of each individual interaction force experienced by the hydrogen atom in fragment 1 due to each atom of non-cysteine amino acids is taken along a vector pointed from H to S. Then the magnitudes of each component was added in order to get the net electrostatic force exerted by the non-cysteine residues of the active site on fragment 1 (hydrogen fragment). The same procedure is repeated for the fragment 2, but the components of individual forces experienced by each atom in fragments 2 due to atoms in non-cysteine residues were calculated along a vector pointed towards S to H. Then the sum of components of these forces is taken to obtain the net force experienced by the fragment 2 due to non-cysteine atoms of the active site. Finally, the forces experienced by the two fragments had been added vectorially to obtain the net force exerted by the external environment of the active site on the S-H bond. The magnitude of all the individual interaction forces was calculated employing the dielectric constant of the water (ε = 78.3553) as per the Coulomb’s Law.

Further, to compute the relative strength of the zwitterionic interaction between Arg207 and Glu21 (see Fig. 2b) in all the four chains of the wt *Ef*BSH and E269A mutant, the electrostatic force of binding is computed along the line joining the central (carbon) atom of the acetate moiety of Glu21 and the central (carbon) atom of the guanidinium moiety of Arg207 as shown in the Figure 2b.

**Fig. 2:**
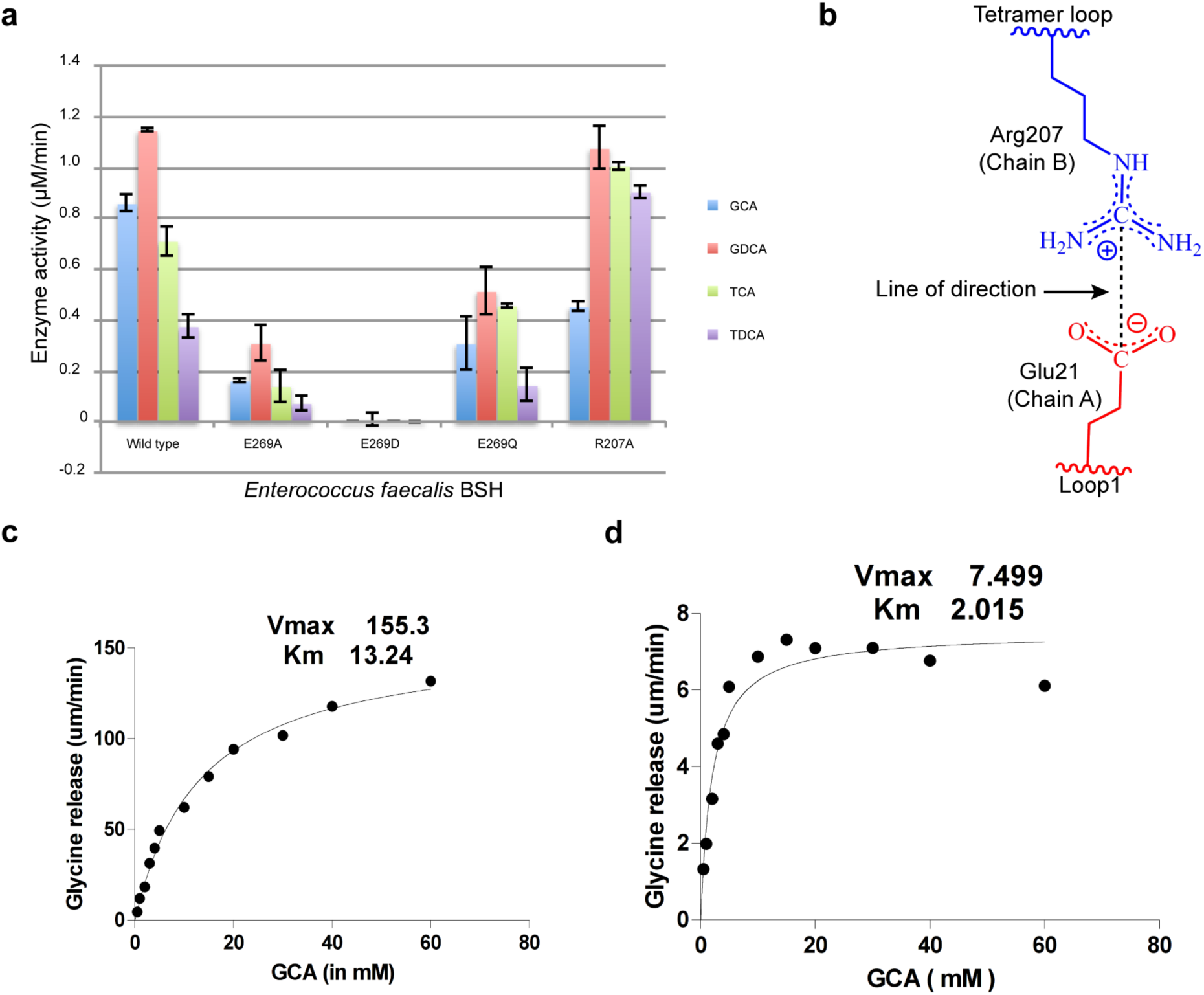
Difference in kinetic and electrostatic behavior of mutant enzymes: **a)** Residual activity profile of *Ef*BSH mutants with GCA, GDCA, TCA and TDCA. **b)** R207A mutant (left) showing shift from allostery to MM kinetics. **c)** Right panel represents E269A mutant showing reduced activity as reflected by Vmax.

The interaction energies for the noncovalent interactions between the acetate group (a side chain analogue of Glutamate) of Glu21and side chain of Tyr20 in the E269A and R207A have been calculated employing the following formula.

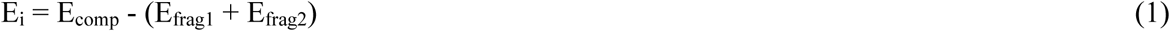

Where, E_comp_ is the energy of noncovalent bonded complex obtained after constrained optimization (as described earlier in the second paragraph of this section), and E_frag1_ and E_frag2_ are single point energies of two interacting fragments being separated from the optimized complex. The starting structure of each complex subjected for the constrained geometry optimization was taken by chopping their structures out from the crystal structures of the respective proteins in such a way that only the noncovalent interactions between side chain of Glu21 and Tyr20 persists in the obtained noncovalent bonded complex. The constrained optimization has been performed to preserve the experimental geometry. The optimized geometries of these noncovalent bonded complexes at the M06-2X (water)/6-31g** level of theory have been shown in Fig. S13 in the supplementary information.

## Results

All the purified proteins (wild type and mutants) were assessed by SEC and were found to be tetramer. These proteins were confirmed using western blot analysis (Fig. S1 a&b).

### Biochemical study

The BSH activity of the selected proteins was carried out by ninhydrin assay to estimate the hydrolyzed by-product, glycine. Percent residual activity of these proteins suggests that the R207A mutant retains the nearly complete *Ef*BSH native activity towards *glyco*-conjugated bile acids (GCA and GDCA), only a marginal decrease in the enzymatic activity was observed in this case. However, an enhanced activity with respect to the wild type *Ef*BSH was observed for the tauro-conjugated bile acids (TCA and TDCA) (Fig. 2a and Table S1). On the other hand, the E269A mutant showed a reduced biochemical activity for both types of substrates (*glyco-* and *tauro*-conjugated bile acids). The E269D mutant was found to be inactive toward both the substrates, while E269Q was found to be active towards them. The enzymatic activity of E269Q is observed to be in between the E269A and the *Ef*BSH (Fig. 2a and Supplementary Information Table 1). Kinetic studies of E269A and R207A mutants revealed the change in kinetic parameters (Fig. 2c & 2d). Significant reduction in V_max_ and enhanced affinity towards GCA substrate (smaller value of K_m_) was observed for E269A mutant. Only minor reduction in the K_m_ value, in comparison to the wild type, was observed in case of R207A. Biochemical studies showed that R207A mutant retains the activity towards GCA and GDCA while it exhibits enhanced activity towards TCA and TDCA (Fig. 2a). The E269A mutant shows reduced biochemical activity towards both *glyco-* and *tauro*-conjugated bile acids. Docking of different conformations of GCA, generated after ligprep, in the active site of *Ef*BSH showed different binding modes (Fig. S11) which restricted the entry of ligand from acceptor site (Chand et al., 2018). The E269D mutant is inactive while E269Q is found to be active towards both *glyco*- and *tauro*-conjugated bile acids. Mutating Glu269 to neutral amino acids i.e. alanine and glutamine retains the activity while reducing the length of side chain by mutating it to aspartic acid completely abolishes the activity towards both *glyco*- and *tauro*-conjugated bile acids.

### Structural Study of *Ef*BSH E269A and R207A mutants

#### Crystallization and data collection

The *Ef*BSH mutant proteins were crystallized using commercial screens by sitting drop method. The good quality diffracting crystals were obtained in 0.1 M Lithium sulfate, 0.1 M Na citrate tribasic dihydrate pH 5.5 and 20% PEG1000. The crystals of mutants R207A and E269A diffracted at 2.0 Å and 1.5 Å respectively. Space group found to be P2_1_ for both the mutants and the unit cell parameters of mutants R207A and E269A are, a = 59.33Å, b = 156.45 Å, c = 72.91 Å; α = γ = 90°, β = 98.62° and a = 59.84 Å, b = 156.32 Å, c = 73.15 Å; α = γ = 90°, β = 98.8° respectively. Matthews’s coefficient confirms the presence of four chains in asymmetric unit in both the mutants. The R207A mutant was refined till R-factor and R-free reaches 20.73 and 23.62, respectively. The E269A mutant was refined till R-factor and R-free reaches 20.35 and 25.18 respectively. Geometry of the residues was further confirmed by Ramachandran plot and found no residues in the disallowed region. Data collection and refinement statistics of both *Ef*BSH mutants is given in table 1. The residues used in the analysis of R207A and E269A mutant were also confirmed by 2Fo-Fc difference maps at 1σ (Fig. S2).

### Structural analysis

We started our analysis with Cys2 (since Met1 is auto-catalytically processed and removed) to map the hydrogen bonding network in the active site pocket of all chains of the *Ef*BSH. The hydrogen bonding pattern near the active site was found to be uniform in all chains for Cys2, Gln256, and Asp19 but varies for Glu269 and Glu21 in different chains (Fig. S3). These variation at the active site resulted from the involvement of Arg207 from other chain of dimer, which protrudes into the active site to interact with Glu21and Glu269 of loop1 and loop4 respectively (Fig. 1c & 3a i-iv). Mutation of Arg207 residue to alanine (R207A) resulted in the release of Glu269 and Glu21 from the hydrogen bonding (Fig. 3b i-iv). However, the mutation of Glu269 to alanine (E269A) resulted into the uniform hydrogen binding interaction between Arg207 and Glu21 in all chains of tetrameric protein (Fig. 3c i-iv).

Structural analysis of Tyr20 upon superimposition of all four chains in the wt *Ef*BSH revealed the scattered arrangement of tyrosine in different chains (Fig S7b). While the similar analyses of Tyr20 of all four superposed chains in the E269A and R207A mutants revealed a congregation of tyrosyl side chains in an identical orientation (Fig. S7 a & c). Moreover, superimposition of four chains of E269A with four chains of R207A showed two extreme conformations possible for Tyr20 in two cases (Fig S8). Interestingly, Tyr20 of the wt *Ef*BSH is found to be drifting between these two extremes (Fig.4).

Structural superposition analysis of wt *Ef*BSH with the previously characterized BSHs structures with respect to the Arg207, Glu21, Glu269 and Tyr20 of wt *Ef*BSH is performed. The results of the analysis are provided in Fig. S9, S10 and Table S3 in the supplementary information file. Tyr20 of *Ef*BSH is replaced by Trp21 in *Bl*BSH, Ile22 in *Cp*BSH and Leu20 in *Ls*BSH (Fig. S10). Surface view of Tyr20 and its corresponding amino acids in other BSHs with conserved asparagine (crucial in anion hole formation) showed blockage of acceptor site in *Bl*BSH and *Ls*BSH (Fig. S10 & S12).

### Computational study

This resulted into the weaker electrostatic interaction (weaker hydrogen bond) between Arg207 and Glu21 in the wt *Ef*BSH. Interestingly, the computed electrostatic force of binding for the zwitterionic complexes (Arg207 and Glu21) of all the chains in E269A mutant are found to be higher (more negative value) as compared to the respective chains in wt *Ef*BSH (Table 2).

In order to decipher the influence of these mutations on the conformational stability of the Tyr20, the interaction energy between the acetate group (side chain analogue of Glutamate) of Glu21 and Tyr20 of E269A and R207A is calculated using DFT for chain A of E269A and R207A mutants. The interaction energy obtained for the two cases is provided in Table 3. The interaction energy between acetate moiety of Glu21 and the side chain of Tyr20 has been found to be smaller in E269A in comparison to that in the R207A by almost 1.5 kcal/mol. This suggests that the acetate in R207A is more strongly bound with Tyr20, and hence holds the structure tighter in this conformation. Further, the noncovalent complex formed between the side chain analogue of Glu21 and the side chain of Tyr20 in R207A mutant is found to be 1.7 kcal/mol lower in energy as compared to E269A mutant (see Table 3). These results are found to be consistent with different theories that are employed for computing the interaction energies (Table 3).

**Table 3:** The interaction energy between acetate moiety of Glu21 and side chain of Tyr20 in E269A and R207A mutant.

| Level of theory | L<sub>1</sub> | L<sub>2</sub> | L<sub>3</sub> | L<sub>4</sub> |
| --- | --- | --- | --- | --- |
| **E269A** | -0.3 | -1.5 | 0.5 | -0.3 |
| **R207A** | -1.4 | -2.7 | -1.1 | -1.8 |
| **E<sub>R207A</sub>-E<sub>E269A</sub>** | -1.7 | -1.7 | -2.5 | -2.2 |

L<sub>1</sub> = M06-2X(water)/6-311++g\*\*
L<sub>2</sub> = M06-2X(water)/6-31g\*\*
L<sub>3</sub> = B3LYP-D2(water)/6-311++g\*\*
L<sub>4</sub> = ωB97XD(water)/6-311++g\*\*
E = electronic energy of the noncovalent complex obtained after constrained optimization

In order to examine the electrostatic basis of the reduced activity of the E269A mutant, electrostatic force analysis has been performed for the E269A and the wt *Ef*BSH. The influence of the non-cysteine amino acids of the active sites of all the four chains on the S-H bond stretching has been evaluated. The detailed procedure is provided in the computational details section. The results of the force analysis are summarized in Table 4, which suggests that the non-cysteine residues in the wt *Ef*BSH stabilize the S-H bond of cysteine, which is contrary to the experimental finding. Since the S-H bond cleavage is one of the key steps in the catalytic cycle of the BSH, any factor which stabilizes S-H bond is expected to increase the reaction barrier. However, S-H bond cleavage is coupled with the N-H bond formation (intramolecular proton transfer reaction between different groups of cysteine) in the transition state structure (Oinonen & Rouvinen, 2000), which may or may not experience the same effect. A complete electrostatic picture evaluating the long range electrostatic influence of non-cystine residues on the turn over frequency of the enzymatic activity of two proteins (wt *Ef*BSH and E269A mutant) is out of the scope of the current study. Moreover, the electrostatic influence of the non-cystine residues in the active sites of each chain on the S-H bond cleavage is marginal as the difference in the electrostatic binding forces for S-H bond of each chain in two cases is very small.The same is also expected for the N-H bond formation (as the S-H bond cleavage is coupled with N-H bond formation in the cysteine) due to the large separation between the cysteine and the other residues present in the active site of these two proteins. This result is further corroborated by comparing the S-H (covalent) bond lengths, which is found to be almost the same for all the chains in the two proteins (see column 4 and 7 of Table 4). Marginally smaller S-H bond length in E269A suggests that the S-H bond is slightly deactivated in E269A mutant, which is in accordance with our force analysis results, as very small positive values of the forces (which pushes hydrogen atom towards the rest part of the cystine) have been obtained for this case (columns 5 and 6, Table 4). These results indicate that the change in electrostatic force due to mutation of Glu269 in *Ef*BSH to Ala269 will have a very marginal influence on the activity of the E269A mutant. Thereby, it indicates the conformational basis of reduced enzymatic activity for E269A mutant.

**Table 4:**
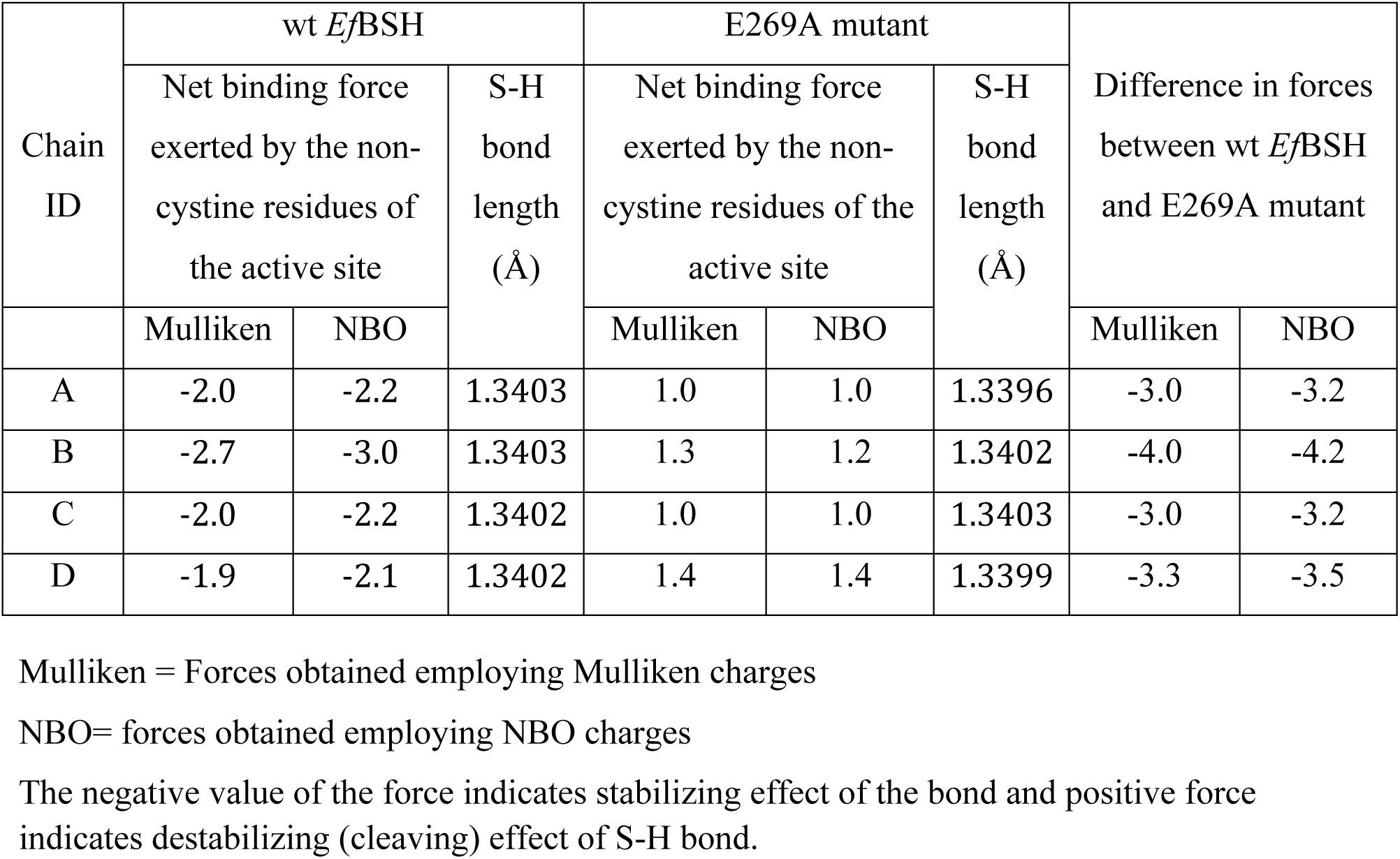
The influence of the non-cystine residues of the active site on the S-H bond cleavage in the pN for the geometries obtained after the constrained optimization at the M06-2X(water)/6-31G** level of theory.

## Discussion

Out of the two mutant *viz. Ef*BSH R207A and *Ef*BSH E269A generated here, E269A mutant significantly loses the biochemical activity towards both glyco- and tauro-conjugated bile acids. The loss in biochemical activity can be attributed either to ionization of S-H bond of catalytic cysteine or it may show conformation basis of reduced activity. In order to examine the electrostatic basis of the reduced activity of the E269A mutant, electrostatic force analysis has been performed for the E269A and the wt *Ef*BSH. The influence of the non-cysteine amino acids of the active sites of all the four chains on the S-H bond stretching has been evaluated. Our result suggests that the non-cysteine residues in the wt *Ef*BSH stabilize the S-H bond of cysteine, which is contrary to the experimental finding. Since the S-H bond cleavage is one of the key steps in the catalytic cycle of the BSH, any factor which stabilizes S-H bond is expected to increase the reaction barrier. However, S-H bond cleavage is coupled with the N-H bond formation (intramolecular proton transfer reaction between different groups of cysteine) in the transition state structure (Oinonen & Rouvinen, 2000), which may or may not experience the same effect. A complete electrostatic picture evaluating the long range electrostatic influence of non-cystine residues on the turnover of the enzymatic activity of two proteins (wt *Ef*BSH and E269A mutant) is out of the scope of the current study. Moreover, the electrostatic influence of the non-cystine residues in the active sites of each chain on the S-H bond cleavage is marginal as the difference in the electrostatic binding forces for S-H bond of each chain in two cases is very small.The same is also expected for the N-H bond formation (as the S-H bond cleavage is coupled with N-H bond formation in the cysteine) due to the large separation between the cysteine and the other residues present in the active site of these two proteins. This result is further corroborated by comparing the S-H (covalent) bond lengths, which is found to be almost the same for all the chains in the two proteins (see column 4 and 7 of Table 4). Marginally smaller S-H bond length in E269A suggests that the S-H bond is slightly deactivated in E269A mutant, which is in accordance with our force analysis results, as very small positive values of the forces (which pushes hydrogen atom towards the rest part of the cystine) have been obtained for this case (columns 5 and 6, Table 4). These results indicate that the change in electrostatic force due to mutation of Glu269 in *Ef*BSH to Ala269 will have a very marginal influence on the activity of the E269A mutant. Thereby, it indicates the conformational basis of reduced enzymatic activity for E269A mutant.

On contrary, *Ef*BSH R207A mutant showed increased biochemical activity towards tauro-conjugated bile acids as compared to glycol-conjugated bile acids. This indicates the involvement of Arg207 in modulation of biochemical activity towards bile acids. Structural analysis showed the position of Tyr20 is locked in two extreme conformations in R207A and E269A mutants, respectively. It has been reported previously that the presence of Tyr20 in loop1 and Tyr65 in loop2 of *Ef*BSH (Fig. 1b) helps in aligning the substrates for the better catalytic framework, and therefore, could be the reason of higher activity. Also loop2 is proposed to transmit the allostery through the tetramer loop (Chand et al., 2018). As predicted previously, we found that Tyr20 from loop1 and Arg207 from tetramer loop are involved in cooperativity. To further explore the pathway of cooperativity, we calculated the interaction energy between the acetate group (side chain analogue of Glutamate) of Glu21 and Tyr20 of E269A and R207A using DFT for chain A of E269A and R207A mutants. The interaction energy between acetate moiety of Glu21 and the side chain of Tyr20 has been found to be smaller in E269A in comparison to that in the R207A by almost 1.5 kcal/mol. This suggests that the acetate in R207A is more strongly bound with Tyr20, and hence holds the structure tighter in this conformation. Further, the noncovalent complex formed between the side chain analogue of Glu21 and the side chain of Tyr20 in R207A mutant is found to be 1.7 kcal/mol lower in energy as compared to E269A mutant (see Table 4). These results have been found to be consistent at different levels of theory that have been employed for computing the interaction energies (Table 4). Although the difference in the energy of the two noncovalent complexes is not very high, nevertheless the relative occupancy of the noncovalent-bonded complex in R207A mutant would be almost 18 times higher than that in the E269A mutant as per the Boltzmann distribution law.

However, the mutation of Glu269 to alanine (E269A) resulted into the uniform hydrogen binding interaction between Arg207 and Glu21 in all chains of tetrameric protein (Fig. 3c i-iv). The interaction between Arg207 and Glu21 in the wild type is expected to have similar strength (and hydrogen bonding pattern) as compared to the E269A mutant. However, the presence of an extra glutamate residue from loop4 (Glu269) near the R207 in the active site of the wt *Ef*BSH (at the place of Ala in the E269A mutant) creates an electrostatic pull (due to its spatial arrangement in the 3D space) to attract the Arg207 towards itself (Fig. S5), and hence increases the distance between Arg207 and Glu21. This resulted into the weaker electrostatic interaction (weaker hydrogen bond) between Arg207 and Glu21 in the wt *Ef*BSH. Interestingly, the computed electrostatic force of binding for the zwitterionic complexes (Arg207 and Glu21) of all the chains in E269A mutant have been found to be higher (more negative value) as compared to the respective chains in wt *Ef*BSH (Table 3). Thus, the active site pocket in the wt *Ef*BSH is more flexible as compared to the E269A mutant and can easily allow to passing the substrate inside the pocket, which is one of the crucial steps involved with the enzyme catalysis. This suggests the electrostatic role of Glu269 in maintaining the dynamics of active site. In case of E269A mutant, Arg207 form triplet of hydrogen bond and forms stronger hydrogen bonding network with Glu21 (Friesner et al., 2006). Also, in the E269A mutant, the electrostatic pull created by the Glu269 gets vanished, and hence resulted into smaller hydrogen bond length (and stronger electrostatic interaction) between Ar207 and Glu21 (Table 3). This leads to the reduction in the dynamics of the active site, which has been further corroborated by the RMSD analysis of the E269A and the wt *Ef*BSH (Fig. S6 a-c).

Structural superposition analysis of wt *Ef*BSH with the previously characterized BSHs structures with respect to the Arg207, Glu21, Glu269 and Tyr20 of wt *Ef*BSH has been performed. The results of the analysis have been provided in Fig. S9, S10 and Table S3 in the supplementary information file. Tyr20 of *Ef*BSH is replaced by Trp21 in *Bl*BSH, Ile22 in *Cp*BSH and Leu20 in *Ls*BSH (Fig. S10). Surface view of Tyr20 and its corresponding amino acids in other BSHs with conserved asparagine (crucial in anion hole formation) showed blockage of acceptor site in *Bl*BSH and *Ls*BSH (Fig. S10 & S12). This is, indeed, supported by our previous studies which mutated the conserved asparagines (Asn79 in *Ef*BSH) to tyrosine and tryptophan. Mutating Asn79 to tryptophan (N79W) drastically reduces the biochemical activity while tyrosine mutant (N79Y) showed marginal decrease in biochemical activity (Chand et al., 2018). Perhaps these residues are complementing each other to form the gate near the active site and the entry of substrate into the active site is directly related to the bulkiness of these residues. As the Asn79 is conserved in all BSHs (Fig. S12), any difference in activity profile of this enzyme is attributed to residues corresponding to Tyr20 in loop1of *Ef*BSH. Hence, Tyr20 in *Ef*BSH acts as a gatekeeper residue.

The BSH activity of the selected proteins was carried out by ninhydrin assay to estimate the hydrolyzed by-product, glycine. Percent residual activity of these proteins suggests that the R207A mutant nearly retains the *Ef*BSH activity towards *glyco*-conjugated bile acids (GCA and GDCA). Only a marginal decrease in the enzymatic activity for this case has been observed. However,an enhanced activity with respect to the wild type *Ef*BSH has been observed for the tauro-conjugated bile acids (TCA and TDCA) (please see Fig. 3a and Table S1 in the Supplementary Information). On the other hand, the E269A mutant showed a reduced biochemical activity for both types of substrates (*glyco-* and *tauro*-conjugated bile acids). The E269D mutant has been found to be inactive toward both the substrates, while E269Q has been found to be active towards them. The enzymatic activity of E269Q has been observed to be in between the E269A and the *Ef*BSH (Fig. 3a and Table S1). Kinetic studies of the E269A and R207A mutant revealed the change in kinetic parameters (Fig. 3c & 3d). A significant reduction in V_max_ and a significantly enhanced affinity towards GCA substrate (smaller value of K_m_) have been observed for E269A mutant. Only a minor reduction in the K_m_ value in comparison to the wild type has been observed for R207A. Biochemical studies showed that the R207A mutant retains the activity towards GCA and GDCA and while showed enhanced activity towards TCA and TDCA (Fig. 3a). Docking of different conformations of GCA, generated after ligprep, in the active site of *Ef*BSH showed different binding mode (Fig. S11) which restricted the entry of ligand from acceptor site (Chand et al., 2018). The E269D mutant is inactive while E269Q is found to be active towards both *glyco*- and *tauro*-conjugated bile acids. Mutating Glu269 to neutral amino acids i.e. alanine and glutamine retains the activity while reducing the length of side chain by mutating it to aspartic acid completely abolishes the activity towards both *glyco*- and *tauro*-conjugated bile acids. This confirms the electrostatic role of Glu269 of loop4 in the active site. The qualitative electrostatic potential surface of E269A mutant showed enhanced electropositive nature of active site. This indicates the role of Glu269 in maintaining the electrostatic nature of active site essential for the catalysis (Fig. S4). It is to be noted that the different electrostatic potential (*viz*. electrostatic field) have been reported to be crucial for the catalysis of different steps of the reaction in the catalytic process of an enzyme (Lai, Chen, Cho, & Shaik, 2010).

**Fig. 3:**
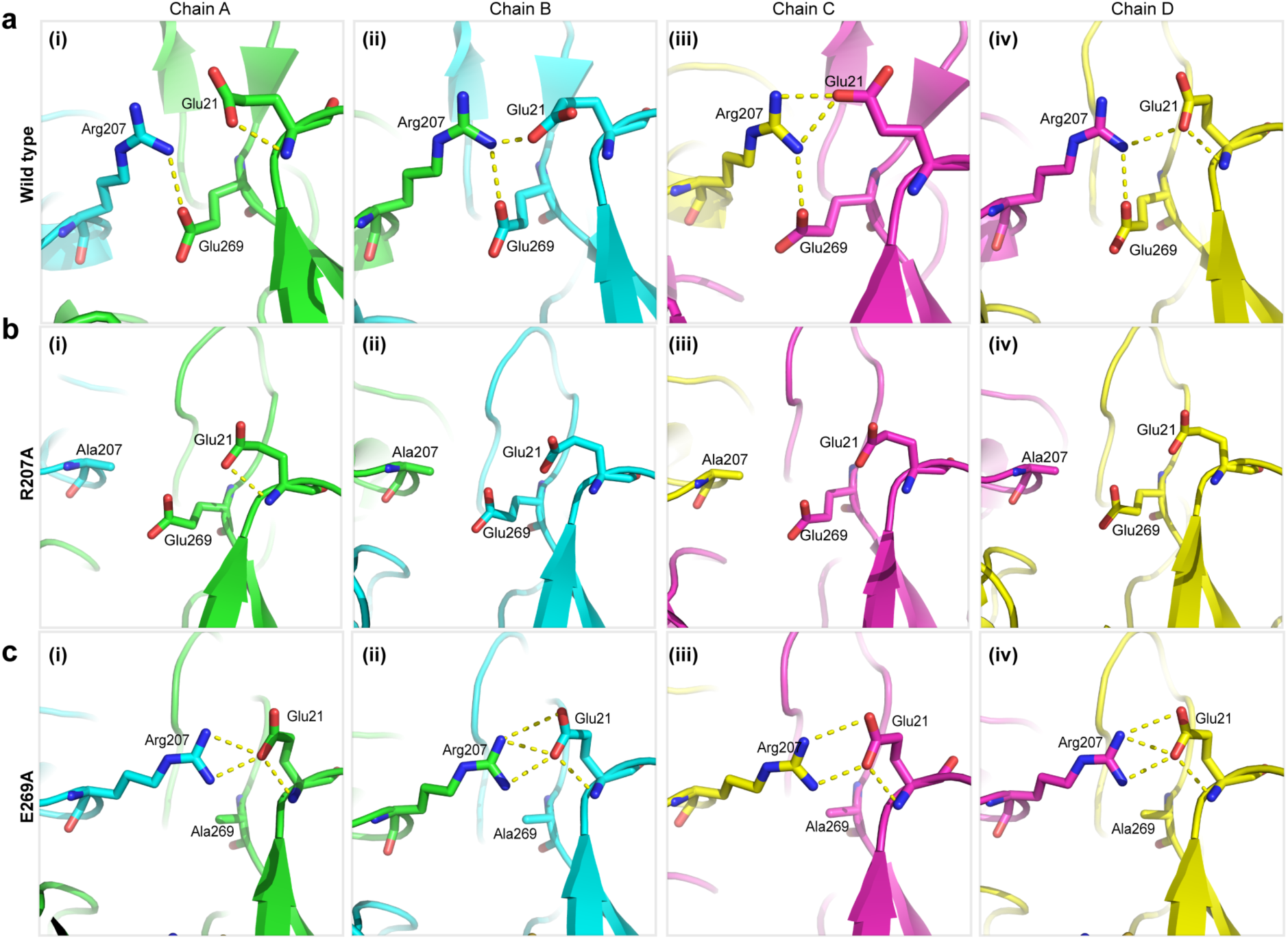
hydrogen bond dynamics of Active site: Top panel describes the dynamics of active site in wild type *Ef*BSH between Arg207, Glu21 and Glu269 (shown in stick). Middle panel showed no interaction between the specified residues after mutating Arg207. Lower Panel describes the E269A mutant which showed that the residues are hydrogen bonded in all the chains.

Thus, it can be concluded that the differential interaction between Arg207 and Glu21 directs the movement of Tyr20 in different forms of *Ef*BSH. In E269A mutant, interaction energy calculation suggests that Glu21 does not interact with Tyr20 and biochemical data showed very less activity as compared to the wild-type *Ef*BSH. Moreover, docking of GCA in active site of *Ef*BSH E269A mutant restricted the entry of substrates from Site A encompassing the loop1 to loop3 (Fig. S11). Therefore this conformation of Tyr20 can be designated as closed conformation. While in case of R207A mutant, the Glu21 showed higher interaction energy with Tyr20 and showed increased activity towards tauro-conjugated bile acids. Therefore, this conformation of Tyr20 can be designated as open conformation (Fig. 4). These results show that without Arg207 the active site is open in all chains and involvement of Arg207 regulates the opening and closing of active site via Glu269 and Glu21. These results are further complemented by biochemical data which showed higher percent residual activity of R207A mutant with tauro-conjugated bile acids (Fig. 3a and Table S1). This is further supported by the fact that previously reported BSH structures have different residues at corresponding location (Fig. S9 a-c) which fails to form either of these combinations (R. S. Kumar et al., 2006; Rossocha et al., 2005).

**Fig. 4:**
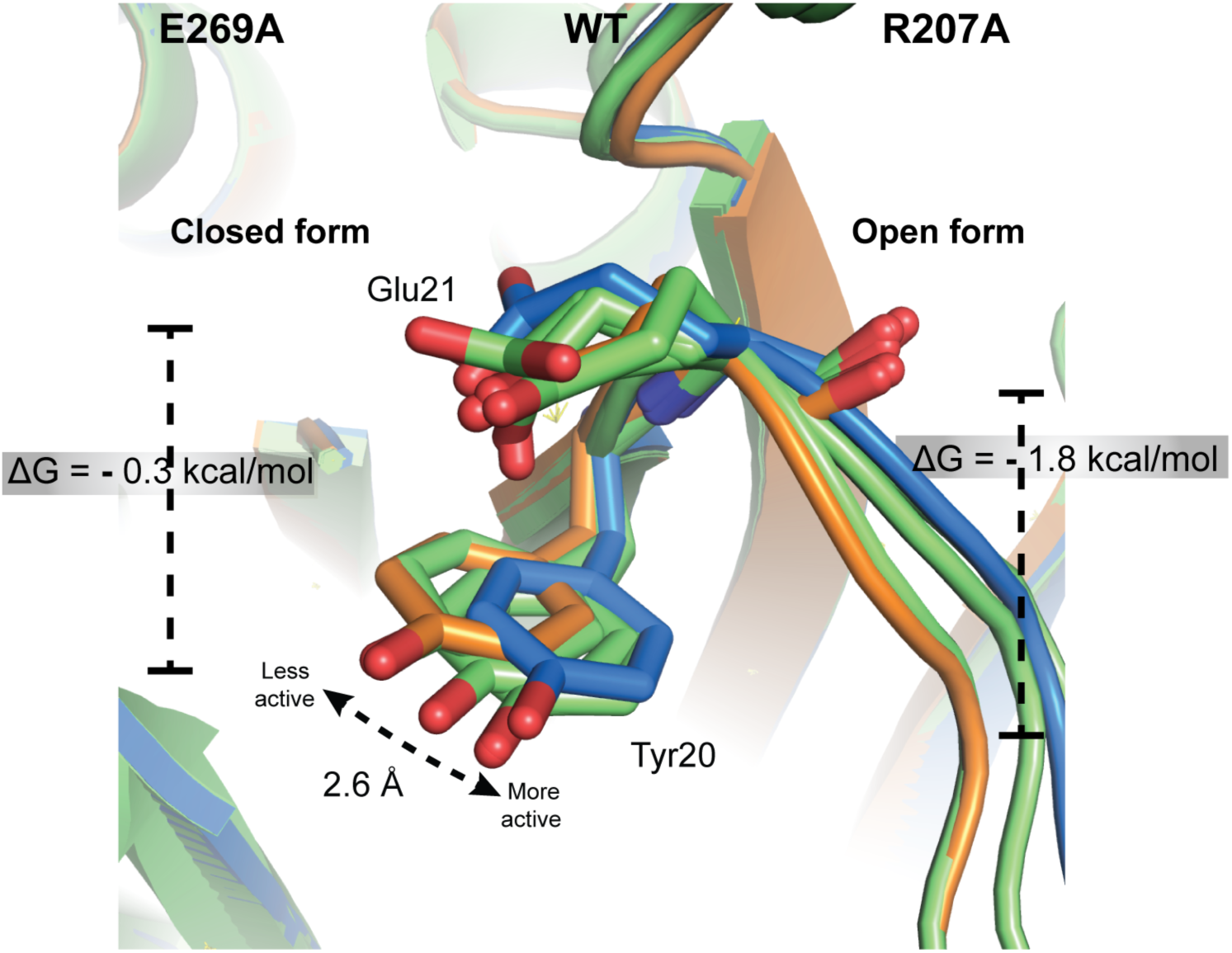
Proposed active site dynamics in *Ef*BSH: Shown here is superposed cartoon image of E269A (orange), WT *Ef*BSH (Green) and R207A (Blue). All four chains of WT *Ef*BSH were used while ChainA of E269A and R207A were used to simplify. Sticks of Glu21 and Tyr20 are shown for all chains.

## Conclusions

In conclusion, we found that the interaction between Arg207 and Glu21 is weaker in nature due to electrostatic pull from Glu269, which results in opening and closing of active site via Tyr20. The outcome of this study would help in engineering of bile salt hydrolases for enhanced deconjugating efficiency towards various bile acids. Further it will be interesting to explore the comprehensive role of Arg207 in the allostery and active site dynamics.

## Acknowledgement

The authors thank beam line scientists of PX-BL21, INDUS-II, RRCAT, Indore for helping in X-ray data collections. The authors also thank the Centre of Excellence in Scientific Computing (COESC), CSIR-NCL, Pune, for providing computationalFacilities.YY thanks University Grant Commission (UGC) for the Senior Research Fellowship. SKR thanks the Department of Science and Technology (DST) for Ramanujan Fellowship and SERB-Young Investigator Grant.

The authors declare no competing interests.

## Coordinate file for Reviewer references

**https://drive.google.com/drive/folders/1svRlSON_xxKATUdzA3MKEPmpKird3ckm?usp=sharing**

